# Rapid generation of marsupial induced pluripotent stem cells suitable for use in next-generation conservation

**DOI:** 10.64898/2026.11.07.765090

**Authors:** Gerard A. Tarulli, Dorthea K. Hansen, Pranav Adithya, Patrick R. S. Tatt, Louize Zheng, Nicola D. Kolaitis, Katarzyna Filimonow, Naomi Tappe, Erin Scase, Ellen Cottingham, Priscilla Jaichander, Savannah J. Hoyt, Gabrielle A. Hartley, Nicole Tillquist, Rachel J. O’Neill, Neil A. Youngson, Benjamin R. Carone, Sara Ord, Andrew J. Pask, Stephen R. Frankenberg

## Abstract

Marsupials account for a disproportionate share of recent and impending mammalian extinctions, that has prompted investment in the development of “next generation” conservation biotechnologies. These include the generation of genetically selected/modified individuals for improved population fitness. Given the inherent limitations of working with threatened species, a flexible vehicle to develop these technologies is through the generation and editing of induced pluripotent stem cells (iPSCs). However, it is deemed necessary to derive transgene-free iPSCs for conservation target species, which can be highly time consuming and expensive. This study describes the generation of iPSCs from the marsupial fat-tailed dunnart (diPSCs) that serves as a model species for next generation conservation initiatives. These diPSCs exhibit features of totipotency when cells are maintained with exogenous reprogramming factors for long periods, until differentiation potential is required.

Neonatal dunnart fibroblasts were reprogrammed using doxycycline-inducible *Oct4*, *Sox2*, *Klf4*, *Lin28* and *Myc*, and the resulting iPSC lines were characterised at the transcriptomic, epigenetic and functional level (trilineage differentiation, embryoid body formation and mouse blastocyst injection assays). Dunnart iPSCs exhibited robust transcriptional and epigenetic activation of core pluripotency factors (*POU5F1, POU5F3, SALL4*) as well as factors associated with the naive and primed states of eutherian pluripotency (*TBX3, KLF2, TFCP2L1, SOX11, DNMT3A*). Dunnart iPSCs readily differentiated in trilineage and embryoid body formation assays, where they expressed markers of all three germ lineages. When injected into 8-cell mouse blastocysts, dunnart iPSCs integrated into both the inner cell mass and trophoblast, reflecting a capacity to contribute to embryonic and extra embryonic lineages during early development. Further, we show that diPSCs can maintain their potential in the presence of exogenous pluripotency factors, until required for differentiation. This robust iPSC generation and maintenance approach is a valuable avenue for the rapid application of next-generation conservation technologies in threatened species.

## Introduction

Species loss has increased in scale and pace in recent decades, and stem cell technologies are at the forefront of next-generation conservation technologies aimed at stemming this tide. Of these technologies, iPSC development holds much promise due to their relative ease of use, providing avenues to both understanding fundamental regulation of pluripotency and in the practical application of stem cells to species rescue. Two Australian marsupials of high conservation value are the northern quoll (*Dasyurus hallucatus)* and the Thylacine (Tasmanian tiger - *Thylacinus cynocephalus*). A close living relative of both species is the fat-tailed dunnart (*Sminthopsis crassicaudata*), which serves as an ideal model and/or surrogate species for marsupial conservation. Marsupials are represented by 335 known extant species within Metatheria, which together with Eutheria (including “placental” mammals) and Prototheria (including monotremes) comprise all living mammals. Marsupials are distinguished from eutherians by their unique mode of reproduction, which involves a brief gestation with a generally non-invasive placenta, followed by a relatively protracted incubation period in an external pouch (“*in marsupio*”). This unique mode of gestation has the potential to place a fundamentally different developmental context against the way pluripotency is traditionally framed, such as in respect to pre- versus post-implantation embryos and substates of pluripotency.

The gene regulatory network controlling pluripotency in embryonic stem cells (ESCs) and induced pluripotent stem cells (iPSCs) has been extensively studied in human and mouse. By comparison, characterisation and establishment of marsupial pluripotent stem cells remains limited. Marsupial iPSCs have been generated in the Tasmanian devil and gray short-tailed opossum^1, 2^. Comparative studies have begun to reveal both conserved and divergent components of the marsupial pluripotency network. For example, orthologs of core pluripotency regulating genes such as *POU5F1* (*OCT4*), *SOX2*, and *NANOG* are expressed during early marsupial development and in pluripotent cell populations^3, 4^. However, marsupials and monotremes also possess an additional paralog of *POU5F1*—*POU5F3*—with the two genes arising from an ancient duplication event and exhibiting distinct expression patterns during early development^5^. In many non-mammalian vertebrates, POU5F3 plays a central role in maintaining pluripotency and regulating zygotic genome activation, whereas in eutherian mammals these functions have largely been subsumed by POU5F1^6^. The retention and developmental expression of *POU5F3* in marsupials raises the intriguing possibility that elements of an ancestral vertebrate pluripotency network persist in this lineage. Investigating pluripotent stem cells from marsupials therefore offers a unique opportunity to dissect the evolutionary plasticity of pluripotency regulatory networks. Of note, expression of the well-established marker of eutherian naïve pluripotency, *ESRRB*, was reported to be absent in Tasmanian devil and platypus iPSCs despite expression of the primary activator of *ESRRB* transcription in eutherian iPSCs, *NANOG*^1, 7^. This finding is profound given the capacity of ESRRB to substitute for NANOG function in pluripotent cells^8^. It has been speculated that *ESRRB* was recruited to the pluripotency network after the divergence of marsupials and eutherians^7^, involved in the acquisition of earlier implantation and more extensive placentation. This highlights that despite the apparent high conservation of core pluripotency networks, there are likely important features unique to the regulation of marsupial pluripotency that are yet to be revealed, and these may serve as hurdles to the application of induced pluripotency in marsupial species conservation.

A major impediment to progress of next-generation conservation for many species is the lack of access to inherent pluripotent populations (i.e. embryonic stem cells). iPSCs have the potential to overcome this limitation, however it can be difficult to achieve stabilised, transgene-free iPSCs from non-model species, limiting both the application of iPSCs in assisted reproduction technologies and investigation of pluripotency networks in each species. Therefore, circumventing the need to generate transgene-free iPSCs in each species would greatly increase the efficiency of technological advancements that assist in species rescue. This study investigated whether high-quality marsupial iPSCs can be generated through continued exposure to exogenous reprogramming factor expression, and that stabilisation in the absence of exogenous factors is not required for use of these iPSCs in downstream differentiation assays and assisted reproduction technology (ART) applications. To this end, iPSCs were generated in a model marsupial—the fat-tailed dunnart—which has high value as a model and surrogate species in conservation biology.

## Materials & Methods

### Fibroblast Generation

Dunnart fibroblasts were isolated from day 3-10 pouch young after removal of the head and internal organs. Tissue was minced using fine scissors until generation of fragments of 2-3 mm^3^. Tissue was cultured on 0.1% gelatin-coated plates until emergence of cell monolayers. Cells were then expanded through passaging using 1 × TrypLE and maintenance of cell density to allow passaging every 3-5 days without allowing cells to become confluent. Cells of passage 3-6 were used for reprogramming. Fibroblasts were maintained in DMEM (Thermo #10566016) with 10% FCS (Thermo #1010047) and 1% antibiotic/antimycotic (Thermo#15240062). All cell culture was performed at 35°C with a 5% CO_2_ and 5% O_2_ atmosphere.

### Transposase Reprogramming

Dunnart fibroblasts were reprogrammed by transfection of plasmids containing: 1) a polycistronic sequence of five mouse reprogramming factors (AddGene #80485^9^) under the control of a tetracycline response element and flanked by piggyBac inverted terminal repeats, 2) a reverse tetracycline-controlled transactivator also flanked by PiggyBac inverted terminal repeats (AddGene #20910), and 3) a piggyBac transposase that is expressed transiently. Transfection was performed using Lipofectamine 3000 (Thermo #L3000001) as per manufacturer’s instructions. At 7-14 days cells were either passaged onto laminin-511 coated plates for colony picking (Merck #CC160) or colonies directly picked into different iPSC medium formulations for further selection and stabilisation on laminin-511. All formulations used the same base medium of 50:50 Neurobasal:Advanced DMEM/F12 (Thermo #21103049 & 12634010) with 1% Glutamax (Thermo # 35050061), 1% antibiotic/antimycotic, 2% B27 supplement (Thermo #17504001), 1% N2 supplement (Thermo #17502001), 0.1 mM beta-mercaptoethanol (Merck #M3148) and 25 ug/mL BSA fraction V (Thermo #15260037). Lines were established in base iPSC medium supplemented with the following factors: 0.5 µM PD0325901 (Stem Cell Technologies # 72184), 10 ng/mL Leukemia inhibitory factor (LIF, Stem Cell Technologies # #78055), 3 ng/mL activin A (Stem Cell Technologies #78132.1) and 2 µM XAV939 (Stem Cell Technologies #72674) (PLAX medium). Lines were transitioned to additional formulations including 5iLAF^10^ containing 1% nonessential amino acids (Thermo #11140050), 50 μg/mL BSA Fraction V, 1 μM PD0325901, 1 μM IM-12 (Merck #SML0084), 0.5 μM SB590885 (Stem Cell Technologies #100-1656)), 1 μM WH-4-023 (Tocris # RDS541310), 10 μM Y-27632 (Stem Cell Technologies # 72304), 20 ng/ml Activin A, 8 ng/ml FGF2 (Miltenyi Biotec), 20 ng/ml LIF and 0.5% knockout serum replacement (Thermo #10828010), 2iLGö medium containing 10 ng/ml human LIF, 250 μM L-ascorbic acid (Merck #AABH9A955F20), 10 µg/ml recombinant human insulin, 1 μM PD0325901, 1 μM CHIR99021 (Stem Cell Technologies #100-1042), 2.5 μM Gö6983 (Stem Cell Technologies # 72462), 10 μM Y-27632, or 10 ng/ml human LIF, 0.5 µM PD0325901 and 0.5 μM CHIR99021^11, 12, 13^, or 2iL. All formulations received 1 µg/mL doxycycline (Sigma #D9891).

### Sendai Reprogramming

Dunnart fibroblasts were treated with CTS CytoTune-iPS 2.1 Reprogramming Kit (Thermo #A16517) as per manufacturer’s instructions with the following modifications. Following Sendai inactivation procedures, cells were passaged at confluence until colonies with pluripotent morphology emerged. At three weeks post transduction colonies were picked, digested in TrypLE for 2 minutes at room temperature and plated into 1 % laminin-511 coated wells (Sigma-Aldrich #CC160). Lines were established in base iPSC medium supplemented with PLAX as above, or LIF, 3 µM CHIR99021 (Stem Cell Technologies #100-1042), 2 µM (*S*)-(+)-dimethindine maleate (Sigma-Aldrich #SML2165-25mg) and 2 µM minocycline hydrochloride (Tocris #3268) - LCDM medium^14^.

### RNA Reprogramming

Dunnart fibroblasts were treated with StemRNA 3^rd^ generation Reprogramming Kit (REPROCELL #00-0076) as per manufacturer’s instructions with the following modifications. Once colonies emerged, cells were bulk passaged onto 1% laminin-511 coated plates (Sigma-Aldrich #CC160) in 50:50 basal iPSC medium (as above):NutriStem hPSC XF (Sartorius #05-100-1A).

### Karyotype analysis

Dunnart iPSCs were passaged 1 to 3 days prior to preparation of chromosome spreads. Metaphase arrest was induced by incubating cells with 0.1 µg/mL KaryoMAX Colcemid solution (Gibco #15212-012) in iPSC culture medium at 35°C for 18 hours depending on the rate of proliferation. Harvested cells were treated with a 0.075 M KCl hypotonic solution that had been preheated to 37°C for 15 min, followed by fixation in methanol/glacial acetic acid (3:1, v/v). Fixed cell suspensions were dropped from approximately 5 cm onto glass slides (Trajan series 2 #472042491). Chromosome spreads were stained by slide immersion into a 0.5 µg/mL DAPI solution (Merck #D1306) for 1 min. Stained slides were covered with glycerol-based fluorescent mounting medium then a coverslip and sealed with clear nail polish. Metaphase spreads were captured and using the Zeiss Axioplan II upright microscope. For male dunnart iPSC lines, metaphase spreads containing 13 morphologically discernible chromosomes were classified as normal, as the extremely small Y chromosome is difficult to reliably resolve.

### RNA Extraction and qPCR

RNA extraction and purification was performed using the GeneElute total RNA purification kit (Sigma-Aldrich #RNB100) as per manufacturer’s instructions. RNA concentration was determined using the Nanodrop One Spectrophotometer (ThermoFisher Scientific). RNA was frozen and stored at −80°C or used in cDNA synthesis using the SuperScript IV First Strand Synthesis System Kit (Invitrogen; #18091050), as per manufacturer’s instructions). Quantification of gene expression was performed by RT-qPCR using the PowerTrack™ SYBR Green Master Mix (Applied Biosystems; A46012) and a QuantStudio 5 Real Time PCR System (Applied Biosystems) in a 10 µL reaction. An initial activation of 95°C for 2 minutes, followed by 40 cycles of 95°C for 5 seconds, and 60°C for 30 seconds was used. Gene- and species-specific primers were designed to amplify regions spanning introns (Supp. Table 1) and all primers presented with a single curve by melt analysis. Primer products were Sanger sequenced to verify specificity. Fold change was determined using a one-way analysis of variance (ANOVA) with Tukey’s post-hoc test. Statistical analysis was performed using GraphPad Prism. Significance was set at a P-value of <0.05 and data is expressed as mean ± SD.

### Bulk RNA Sequencing, Differential Gene Expression & Gene Ontology Analysis

RNA quality was assessed using the High Sensitivity RNA ScreenTape (Agilent) on the 4150 TapeStation (Agilent) as per the manufacturer’s instruction. RNA libraries were prepared using the Illumina Stranded mRNA Prep Kit and sequenced using the Illumina NovaSeq 6000 to generate 50 million 100bp paired-end reads per sample. All cDNA library preparation and sequencing was performed at the University of Connecticut (USA). Raw sequencing reads were assessed for quality using FastQC (v0.12.1) and summarised with MultiQC (v1.25.2). Adapter sequences and low-quality bases were timed using Trimmomatic (v0.39). Reads were then aligned to the Sminthopsis crassicaudata reference genome (NCBI RefSeq assembly GCF_048593235.1) using Kallisto (v0.51.1) with default parameters. Raw count matrices were imported into R (v4.5.1). Genes with fewer than 1 count in at least 3 samples were excluded. Library sizes were normalised using the trimmed mean of M-values (TMM) method and converted to CPM with log2 transformation. Differential expression between culture conditions was assessed using edgeR (v4.0.5) and limma (v3.58.1). Differentially expressed genes were defined using an adjusted P value < 0.01 and an absolute log2 fold change ≥ 2 for downstream DEG selection. Gene ontology analysis was performed using gProfiler against human ontologies.

### Immunofluorescence

Cells were grown on coverslip glass chamber slides (Ibidi #80826) and fixed with 4% paraformaldehyde for 15 minutes at room temperature after which samples were washed with PBS for 5 minutes at room temperature. Immunolabelling was performed by diluting primary antibodies in PBS with 10% normal donkey serum (Merck #D9663) and incubation overnight at 4 °C (Supp. Table 2). After labelling samples were washed with PBS twice for 5 minutes. At this time, secondary antibodies were diluted in PBS with 10% normal donkey serum (Merck #D9663) and incubated at room temperature for 1 hour. Subsequently, samples were incubated with 500 nM DAPI for 5 minutes to label nuclei, after which samples were washed twice for 5 minutes with PBS. Samples were left in the final PBS wash for imaging using a Nikon A1R confocal microscope under sequential scan mode with a z-resolution of 1 Airy unit.

### Trilineage Differentiation

Differentiation of diPSCs into the three primary germ layers was performed using a commercially available kit designed for human iPSCs (STEMdiff Trilineage Differentiation Kit, STEMCELL Technologies #05230) with minor modifications to the manufacturer’s protocol. Briefly, 5 x 10^4^ diPSCs were seeded on 12-well plates coated with laminin-511 (Merck #CC160) and cultured with iPSC medium (with doxycycline) for 48 hours. At this timepoint, 50% of iPSC medium was replaced with trilineage differentiation medium for each lineage test. Control wells had 50% of iPSC medium refreshed. After another 24 hours, a complete medium change was performed on each well that was repeated every 24 hours for an additional 2 days. Cells were harvested for RNA analysis using an RNeasy mini kit (Qiagen #71204), by lysing cells directly in wells and extracting RNA as per manufacturer’s protocol. For immunostaining, samples were fixed and immunostained 48-hours after transition to differentiation medium.

### Embryoid Body (EB) Formation Assay

On day of EB formation, 1 x 10^5^ diPSCs were placed in 6-well ultra-low adherence culture plates (Corning #3471) in diPSC growth medium supplemented with 4 mg/mL polyvinyl alcohol (Sigma-Aldrich #341584) and ROCK pathway inhibitor (Stem Cell Technologies #72378) to assist EB formation and incubated at 35 °C for 24 hours. Medium was then changed to diPSC medium without ROCK inhibitor and PVA and cells were cultured for an additional 48 hours. Subsequently, medium was replaced with EB culture medium (DMEM/F12 supplemented with 10% fetal calf serum (Gibco #72308), 1% sodium-pyruvate (Gibco #11360070), 1% GlutaMAX (Gibco #35050061) and 1% antibiotic/antimycotic (Gibco #15240062) and EBs were cultured for an additional 9 days with daily medium changes. At this time, EBs were transferred to 8-well chamber slides (Ibidi #80826) by embedding in 50% growth factor-reduced matrigel (Corning #356231) diluted in EB culture medium. EBs were left overnight before addition of 200 uL of EB culture medium containing 1% matrigel. EBs were subsequently incubated for an additional 18-21 days with medium changes every 2 days. At the end of experimentation EBs were fixed with 4% paraformaldehyde for 15 minutes, washed with PBS for 5 minutes and kept at 4 °C for immunolabelling. Immunolabelling of EBs was performed in 8-well chamber slides while samples were embedded in matrigel. Samples were first washed with PBS (1 x 5 minutes) and permeabilized in 0.05% TritonX-100 in PBS, overnight at 4 °C. EBs were subsequently immunolabelled with antibodies directed to GATA2 and GATA6 diluted in PBS containing 10% normal donkey serum (Merck #D9663) at dilutions outlined in Supp. Table 1, overnight at 4 °C. The following day, samples were washed twice for 15 minutes in PBS, followed by the addition of fluorescent label-conjugated secondary antibodies diluted in PBS with 10% normal donkey serum for 2 hours at room temperature. This was followed by twice washing samples in PBS for 15 minutes, after which sample nuclei were stained with 500 ng/mL DAPI (Merck D1306) for 5 minutes at room temperature. DAPI was removed from samples and replaced with PBS for imaging. Samples were imaged using a Nikon A1R confocal microscope under sequential scan mode at <1 Airy unit.

### CUT&RUN

CUT&RUN was performed to assess H3K4me3, H3K27me3, H3K9me3, H3K27ac, and CTCF using 100,000–150,000 cells per condition in triplicate (n = 3), following the manufacturer’s instructions (Cell Signaling #86652). Antibodies were used at 1:50: tri-methyl-histone H3 (Lys4) (CD2D8; Cell Signaling #9751), histone H3 (acetyl K27) (ab4729), tri-methyl-histone H3 (Lys27) (C36B11; Cell Signaling #9733), tri-methyl-histone H3 (Lys9) (D4W1U; Cell Signaling #13969), and CTCF (D31H2; Cell Signaling #3418). Rabbit (DA1E) monoclonal IgG XP® isotype control (Cell Signaling #66362) was used at 1:20 as a negative control. The cells were washed using 1x wash bufffer (1x spermidine, 1x protease inhibitor cocktail) and resuspended in 10ul of conanacavlin A beads per reaction Ten microliters of concanavalin bead suspension were added to washed cells, which were then resuspended in antibody binding buffer (1x spermidine, 1x protease inhibitor cocktail, digitonin solution, nuclease-free water). Target antibodies were added at the indicated dilutions and incubated overnight at 4°C with rotation. Samples were washed with 1 ml digitonin buffer (1x wash buffer, 1x spermidine, 21x protease inhibitor cocktail, digitonin solution, and nuclease-free water), followed by addition of 50 µl pAG-MNase premix (digitonin buffer, pAG-MNASE enzyme) and incubation at 4°C for 1 hour with rotation. Samples were washed twice, resuspended in ice-cold calcium chloride, and incubated at 4°C for 30 min. Reactions were stopped with 150 µl 1x stop buffer at 37°C for 10 min. DNA fragments were isolated using the spin-column method with the DNA purification, binding, and elution buffers supplied in the extraction kit (Cell Signaling #14209). Fragment size and quantity were assessed using a High Sensitivity D5000 TapeStation (Agilent Technologies) before sequencing library preparation. Libraries were prepared using the NEBNext Ultra II DNA Library Prep Kit and sequenced on an Illumina NovaSeq SP flow cell to generate 250-bp paired-end reads.

### CUT&RUN analyses

FASTQ adapter trimming was performed using Trimmomatic v0.40, with QC conducted using FastQC v0.12.1 and MultiQC v1.21. Trimmed reads were mapped to the Sminthopsis crassicaudata reference genome (GCF_048593235.1) using Bowtie2 v2.5.5, and alignments were sorted and indexed using SAMtools v1.22.1. Read distribution was assessed with deepTools v3.5.5 bamCoverage using RPGC (Reads Per Genomic Content) normalization to generate BigWig files. BigWigs were normalized against IgG using bigwigCompare with the --operation subtract function, and IgG-subtracted profiles were visualized at key genomic regions using IGV v2.16.0. IgG-subtracted BigWig files were averaged across replicates using bigwigAverage and converted to bedGraph using the UCSC bigWigToBedGraph function. Peak calling was performed with MACS2 v2.2.7.1 bdgpeakcall for narrow H3K4me3 and H3K27ac peaks and bdgbroadcall for H3K27me3. Differential peak enrichment between fibroblasts and iPSCs was assessed using DiffBind in RStudio. Reads within peak regions were counted using dba.count, with summits = 200 for narrow H3K4me3 and H3K27ac peaks and summits = 500 for broad H3K27me3 regions ^15, 16^. Counts were normalized for library size using dba.normalize, and differential enrichment was assessed using dba.analyze. Sequences from significantly enriched iPSC-specific active histone peaks (FDR < 0.05) were analyzed for motif enrichment using MEME-ChIP (MEME Suite v5.5.9), with JASPAR vertebrate and UniPROBE Mouse motif databases. Differential motif enrichment was assessed using AME (MEME Suite), using iPSC-specific active chromatin peak sequences as the primary set and fibroblast active chromatin peaks as the control.

### Mouse Embryo Injection & Immunostaining

8-cell embryos were obtained from 4 to 8 weeks old F1 (Bl/6 × CBA) female mice superovulated with 5 IU of pregnant mare’s serum gonadotropin (Focus Biosciences # HOR-272) followed after 48 h with 5 IU of human chorionic gonadotropin (Focus Biosciences # HOR-0250), and mated with F1 males. Females with vaginal plugs were autopsied 63–64 h after hCG injection. Embryos at the non-compacted 8-cell stage were collected by flushing oviducts and uteri with GMOPS Plus medium (VitroLife # 26083). Embryos were incubated in the same medium prior microinjection. If embryos started compaction before cell injection, they were incubated for 5-10 min in M2 medium without Ca^2+^ and Mg^2+^ supplemented with bovine serum albumin (BSA; Sigma-Aldrich # A8806)^17^. iPSCs were lifted from substrate with prewarmed (37°C) TrypLE for 3 minutes. Suspended cells were transferred to centrifuge tube and diluted with 2ml of culture media and centrifuged for 5 min (180 rcf). Pelleted cells were then resuspended in ∼200 µL of room temperature iPSCs culture medium (PLAX) with 1% HEPES (Thermo #15630080). Cells were kept at 4°C until injection. Eight iPSCs were injected into the mouse non-compacted 8-cell stage embryo using 12-15 µm inner diameter blunt glass pipette (BioMedical Instruments). Pipette was inserted in between blastomeres after a hole in zona pellucida was made using laser. After injection, embryos were cultured for 72 h (late blastocyst stage) in Advanced KSOM medium (Sigma-Aldrich #MR-101-D). Embryos were fixed in 4% paraformaldehyde (PFA) in PBS with 0.1% Tween 20 (Sigma) and 0.01% Triton X-100 (Sigma) for 20 min in RT. Before immunostaining, permeabilisation in 0.55% Triton X-100 in PBS for 15 minutes and blocked in 10% fetal bovine serum in PBS for 40min were performed, followed by 75 min incubation in primary antibody solution. Primary antibodies used were anti-SOX2 and anti-GATA2 (Supp. Table 2). Secondary Alexa Fluor (Invitrogen)-conjugated antibodies were used at a dilution of 1/400. DNA was visualized using Hoechst 33342 (5 μg/ml, Molecular Probes).

## Results

### Derivation of dunnart iPSCs via multiple reprogramming approaches

With the goal of identifying a flexible and robust reprogramming method for marsupial species with high conservation value, dunnart fibroblasts were reprogrammed using three approaches: 1) Sendai virus-based delivery of human *POU5F1, SOX2, KLF4* and *MYC* (CytoTune™ 2.0), 2) RNA-based delivery of human *POU5F1, SOX2, KLF4, LIN28A* and *MYC* (StemRNA™ 3rd gen.), or 3) piggyBac transposase-induced random integration of tetracycline-inducible mouse *Pou5f1, Sox2, Klf4, Lin28A and Myc* (C5). Within 2 weeks of delivery of reprogramming factors via any approach, compact colonies containing cells with large nucleus:cytoplasm ratio and prominent nucleoli began to emerge (Fig. 1A). For each reprogramming approach, colonies were picked and expanded on several iPSC media established in eutherians and assessed for their ability to maintain this morphology over passage. Using Sendai virus, stable lines were generated using multiple medium formulations, including 2iL and LCDM (Supp. Fig. 1A-B), however transcriptomic analysis showed a persistence of viral RNA for up to 35 passages (Supp. Fig. 1C), rendering the lines unsuitable for functional application.

**Figure 1.**
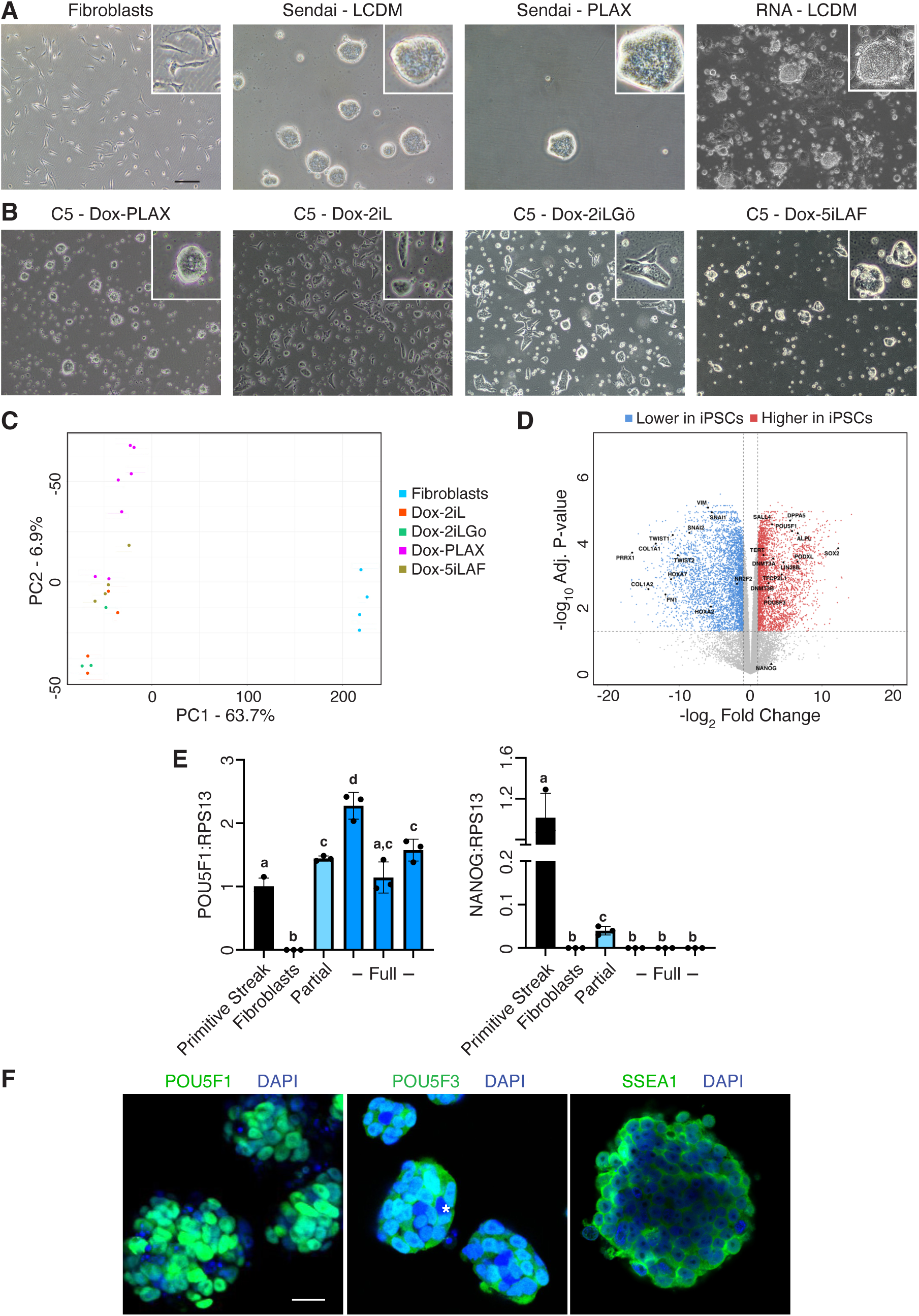
Generation of dunnart iPSCs with stable pluripotency profiles A) Examples of dunnart fibroblasts and iPSCs generated using several reprogramming approaches (Bar = 100 µm). B) diPSCs generated using a randomly-integrated and doxycycline-inducible system cultured under different naive media with doxycycline, showing more consistent morphology with the PLAX (MEKi,WNTi, LIF, Activin) formulation. C) PCA of RNAseq data comparing lines established in different naïve media, showing little medium-specific clustering. D) Volcano plot of fibroblasts versus diPSCs cultured in dox-PLAX highlighting increased expression of core pluripotency and associated factors, along with repression of selected somatic genes. E) qPCR analysis of POU5F1 and NANOG expression in diPSCs during reprogramming and after stabilisation, showing detectable NANOG expression in partially-reprogrammed dunnart iPSCs. *P* F) Dunnart iPSCs cultured in dox-PLAX immunostained with a marsupial-specific POU5F1 and POU5F3 antibodies, and SSEA1. (Bar = 25 µm).

RNA-based delivery offers an attractive alternative to overcome the limitations of persistent viral transgenes, however lines could only be generated that exhibited suboptimal morphology and poor cell health that further degraded over passage (Supp. Fig. 2). By contrast, use of a doxycycline-inducible C5 reprogramming plasmid generated stable lines in multiple media while in the presence of exogenous reprogramming factors (Fig. 1B). These lines maintained a consistent pluripotent morphology over multiple passages, however in accordance with the RNA-based reprogramming method, removal of doxycycline under any culture condition failed to generate stable lines with features of pluripotency. This suggests that either dunnart fibroblasts are either only partially reprogrammed with the C5 method or that reprogrammed cells require unidentified additional culture conditions or reprogramming factors to stabilise the marsupial pluripotency network. Therefore, diPSC could only be generated and maintained with doxycycline-induced exogenous reprogramming factor expression. Nonetheless, the C5 plasmid system is flexible and cost-effective and in the context of species rescue has potential value for the advancement of emerging conservation technologies where knowledge, material and/or financial resources are limited. Therefore, it was hypothesised that using the C5 approach and maintaining diPSCs in the presence of continued exogenous reprogramming factor expression provides a sufficient tool to enable the exploration of next generation conservation technologies in conservation-relevant species. To this end, multiple diPSC lines were derived using the C5 reprogramming approach and subjected to transcriptomic, epigenomic and functional analyses.

**Figure 2.**
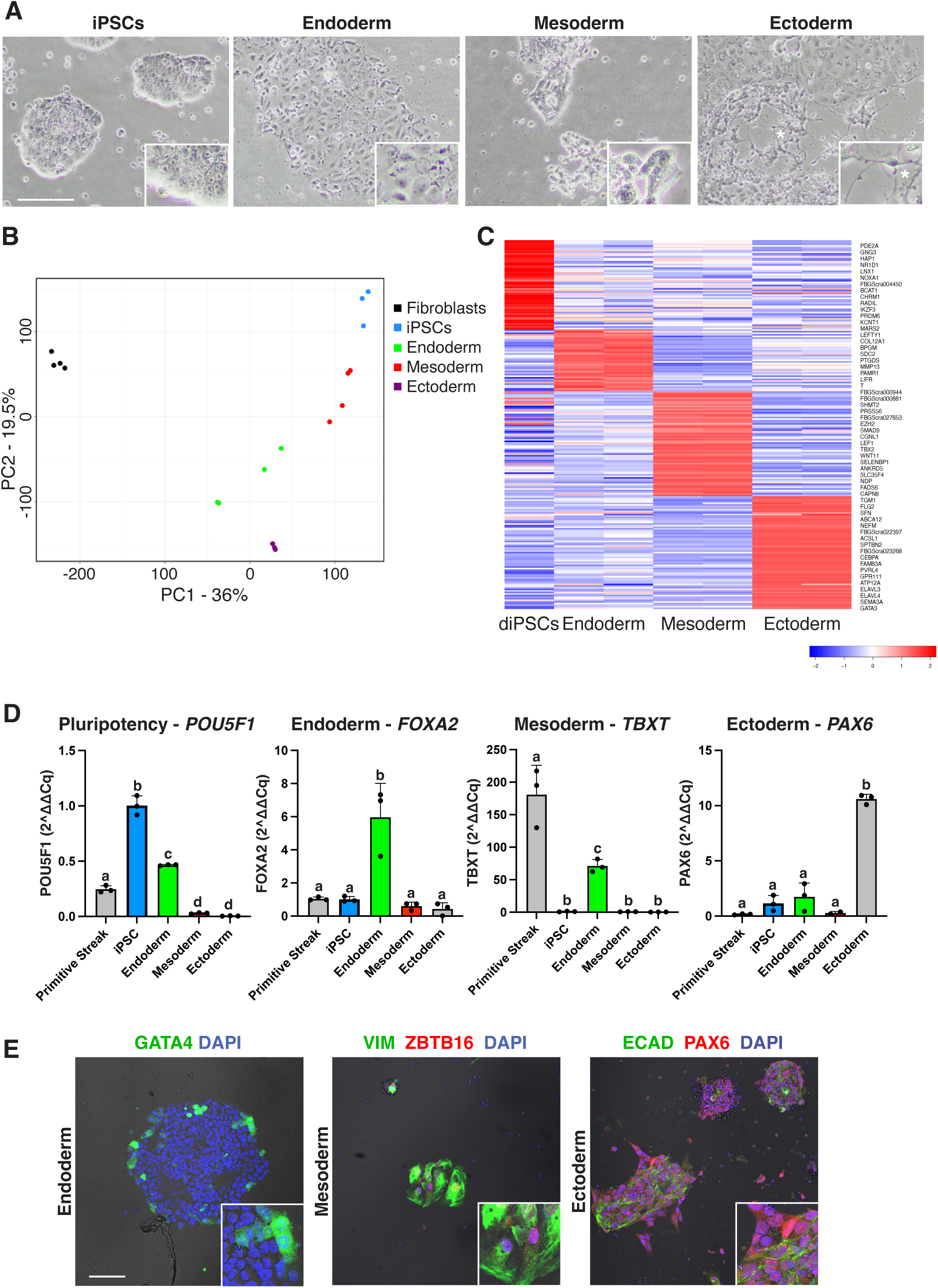
Dunnart iPSCs exhibit robust trilineage differentiation after removal of exogenous reprogramming factors A) Changes in cell morphology four days after removal of doxycycline and induction of trilineage differentiation. B) PCA of two independent dunnart iPSC lines, matched trilineage differentiation samples and fibroblasts showing clustering based on trilineage medium. C) Heatmap of highly expressed genes able to distinguish lineage differentiation in dunnart iPSCs. Expression of many eutherian lineage-specific genes are induced, along with a strong mesendoderm profile in endoderm-differentiated samples. D) qPCR analysis of pluripotency and germ layer differentiation markers before and after trilineage differentiation. Note expression of TBXT in endoederm differentiated samples that reflects a potential mesendodermal differentiation profile. E) Immunofluorescence of proteins associated with germ layer differentiation. (Bar = 100 µm).

Continued expression of reprogramming factors, particularly c-MYC, is associated with genomic instability^18, 19^. Therefore, karyotyping analysis was performed to determine whether a normal karyotype could be retained in diPSCs over a sufficiently high passage number to be suitable for downstream applications. Dasyurids such as the fat-tailed dunnart have a small number of large autosomes^20^. The sex chromosomes are relatively small, and the Y chromosome can be difficult to discern using routine karyotyping methods. Karyotyping was performed on two diPSC lines (Supp. Fig 3A)—one at passage 32 and the second at passage 35, considered a high passage number for iPSCs. In total, 12.7% of cells showed alterations, with 86% of those changes being a lack of a single chromosome (Supp. Fig 3B), with only one cell showing a significant abnormality (tetraploid) (Supp. Fig. 3C).

**Figure 3.**
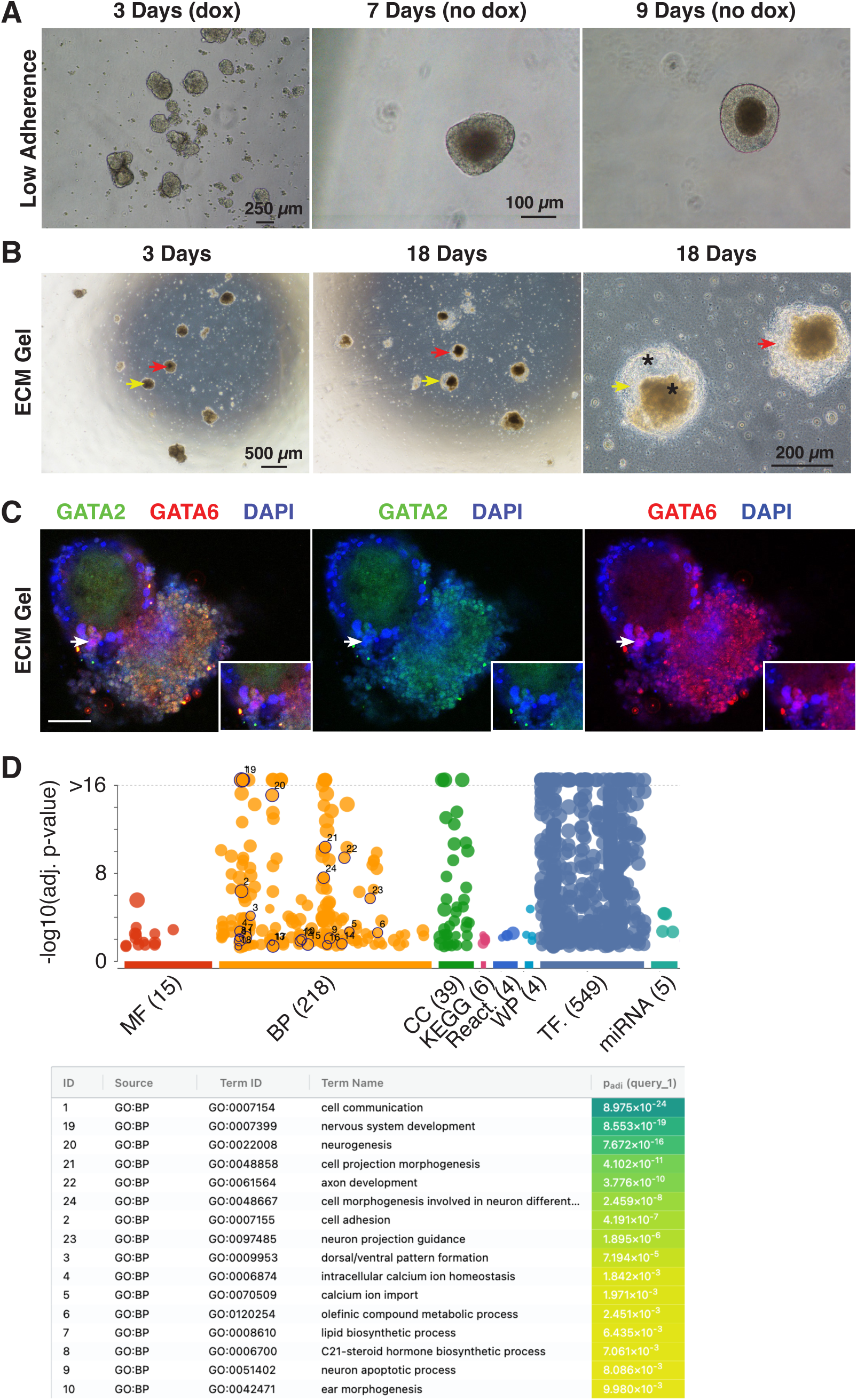
Dunnart iPSCs exhibit organisation and differential expression of trophectoderm and primitive endoderm markers in embryoid body formation assays A) Aggregation and organisation of dunnart iPSCs under low adherence conditions to form embryoid bodies. B) Embryoid body structures after embedding in ECM gel for indicated time periods in the absence of doxycycline. C) Immunofluorescence images of embryoid bodies. Cell expressing GATA6 (red) but not GATA2 (green) indicated by arrow. D) Gene ontology analysis of top 500 genes significantly upregulated in EBs compared to iPSCs. (ULA = ultra-low adherence; Gel = ECM gel matrix).

### Stable diPSCs express many core pluripotency and associated factors but not NANOG and ESRRB in the presence of exogenous reprogramming factors

To determine whether continued expression of reprogramming factor transgenes interferes with the expression of endogenous pluripotency factors in diPSCs, RNAseq analysis was performed under multiple culture conditions. When assessed by principal component analysis (PCA), all iPSC culture conditions occupied a distinct space compared to fibroblasts based on PC1 (63.7%), while different iPSC culture conditions were segregated to a much smaller extent based on PC2 (6.9%) (Fig. 1C). This relatively slim differential suggests that continued exogenous transgene expression does not result in highly distinct transcriptomic profiles of diPSCs when maintained in different culture conditions. Due to the superior morphology of diPSCs grown in PLAX, we selected this medium formulation for more detailed study. Comparative transcriptomic analysis comparing fibroblasts with diPSCs showed strong increases in well-recognised pluripotency-associated genes, including *POU5F1*, *TERT, DNMT3A/B* and the *POU5F1* paralogue, *POU5F3* (Fig. 1D and Supp. Table 3), in concert with reductions in somatic genes including members of the *HOX*, *SNAI* and *COL* families. Further, diPSCs were assessed for transcription of 39 genes upregulated in published studies of mouse, pig, chicken and monotreme iPSCs when compared with respective fibroblasts, and significant upregulation of these genes was also identified in diPSCs (Supp. Fig 4). A notable absence from the transcriptional profile of diPSCs was the core pluripotency factor, *NANOG,* and its downstream target gene, *ESRRB*, in stable diPSC lines. This was in spite of high expression of the eutherian upstream activator of *NANOG SALL4*, and apparent high LIF/STAT3 activity as determined by high levels of *GBX2*, *PIM1* and *SOCS3*^21^. Due to the otherwise strong pluripotency profile detectable by RNAseq, it was speculated that since *NANOG* is known to have its most fundamental role in the acquisition of pluripotency in eutherians (Silva et al 2009), it may be expressed only during the maturation phase of diPSC formation. To test this, RNA was extracted from partially or fully reprogrammed diPSCs and assessed for *POU5F1* and *NANOG* by qPCR (Fig 1E). While *POU5F1* was robustly expressed in partially and fully reprogrammed diPSCs, *NANOG* was only detectable in partially reprogrammed diPSCs. To verify that pluripotency-associated factors are expressed at the protein level, immunofluorescence was performed on fixed diPSCs while exposed to exogenous reprogramming factors. Using antibodies raised to marsupial POU5F1 and POU5F3^3^, clear expression of both key pluripotency factors in diPSCs was observed, with POU5F1 restricted to the nucleus of all live cells to varying degrees and POU5F3 present in both the cytoplasm and nucleus of the majority of cells, with only a small number of diPSCs being POU5F3-negative (Fig 1F, asterisk).

**Figure 4.**
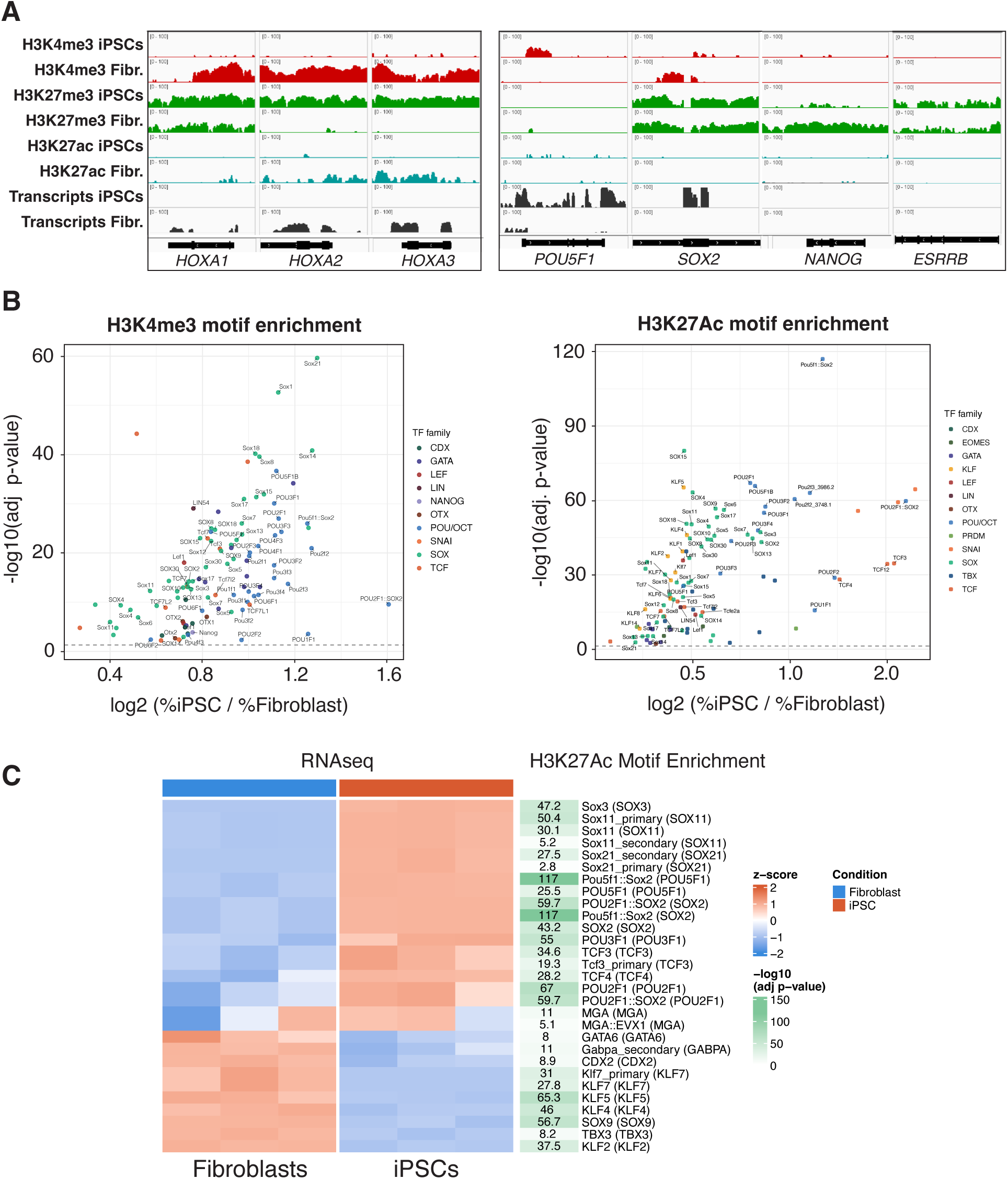
Dunnart iPSCs have epigenetic signatures strongly associated with pluripotency and early embryo development A) Traces of CUT&RUN outputs for selected somatic (left) and pluripotency genes (right), showing repression of somatic genes and activation of a subset of many key pluripotency genes. B) Motif enrichment analysis of H3K4me3 and H3K27Ac marks in dunnart iPSCs versus fibroblasts. Key categories of pluripotency and general differentiation gene families are highlighted. The strongest motif enrichment is for *POU5F1*, *POU2F1* and *SOX2*. C) Association of H3K27Ac-enriched motifs with transcript levels in dunnart iPSCs. H3K27Ac motifs are enriched for a subset of actively transcribed pluripotency genes.

### diPSCs express genes associated with all germ layers after removal of exogenous reprogramming factors and trilineage differentiation

The PLAX medium selected for further study was used to functionally test the hypothesis that sustained exposure of diPSCs to ectopic reprogramming factors does not hinder normal differentiation, thus permitting their application to assisted reproduction technologies. First, diPSCs were tested for their capacity to form cells of the three primary germ layers: endoderm, mesoderm and ectoderm using a modified commercial human trilineage differentiation protocol. Within 4 days of doxycycline removal and transition to lineage-specific medium, diPSCs showed distinct morphological changes associated with each lineage (Fig. 2A), including axonal-like projections in ectoderm-differentiated diPSCs. This assay was performed on two independent C5/PLAX lines, and samples collected for RNA sequencing. PCA analysis showed strong clustering of iPSCs distinct from trilineage differentiated samples and fibroblasts (Fig. 2B), with clear germ layer-specific clustering of differentiation treatments. Differential expression analysis identified many genes with known roles in each lineage being most highly expressed in the respective trilineage sample. This included *FOXA2*, *PTGDS* and *LEFTY1* in endoderm-differentiated diPSCs, *LEF1*, *CDX1* and *AXIN2* in mesoderm-differentiated diPSCs, and *GATA3*, *TUBB3* and *DSC1* in ectoderm-differentiated diPSCs (Fig. 2C & Supp. Table 4). Of note, the classic mesoderm markers *TBXT* and *EOMES* were most highly expressed in endoderm-differentiated samples. For TBXT, this was confirmed by qPCR (Fig. 2D). It was hypothesised that due to the abridged nature of the modified trilineage differentiation protocol, endoderm samples were exhibiting a mesendoderm phenotype^22, 23^. Indeed, when the differentiation protocol was modified to a longer culture duration, *TBXT* was most highly expressed in mesoderm-differentiated iPSCs (Supp. Fig. 5A). Genes that were both highly expressed and specific to each group were selected for gene ontology analysis (Supp. Fig 6). Mesoderm-differentiated samples showed enrichment for mesoderm-associated ontologies, including vascular and skeletal development, while ectoderm-differentiated samples were enriched for epidermal and neuronal ontologies. Endoderm-differentiated samples were enriched for genes associated with collagen and growth factor binding and cartilage development, as well as general processes including multicellular organism development, however no significant enrichment of endoderm-specific ontologies was identified. Nonetheless, significant upregulation of primitive endoderm markers was observed in endoderm-differentiated diPSCs, including *GATA6* and *PDGFRA*. Expression of lineage-associated genes was also confirmed at the protein level, with endodermal GATA4, mesodermal PLZF and ectodermal PAX6 expression detected in the respective lineages (Fig 2E). Together, our data suggest that prior maintenance of continued exogenous reprogramming factors expression does not inhibit the capacity of diPSCs to form three primary germ layers in a trilineage differentiation assay.

**Figure 5.**
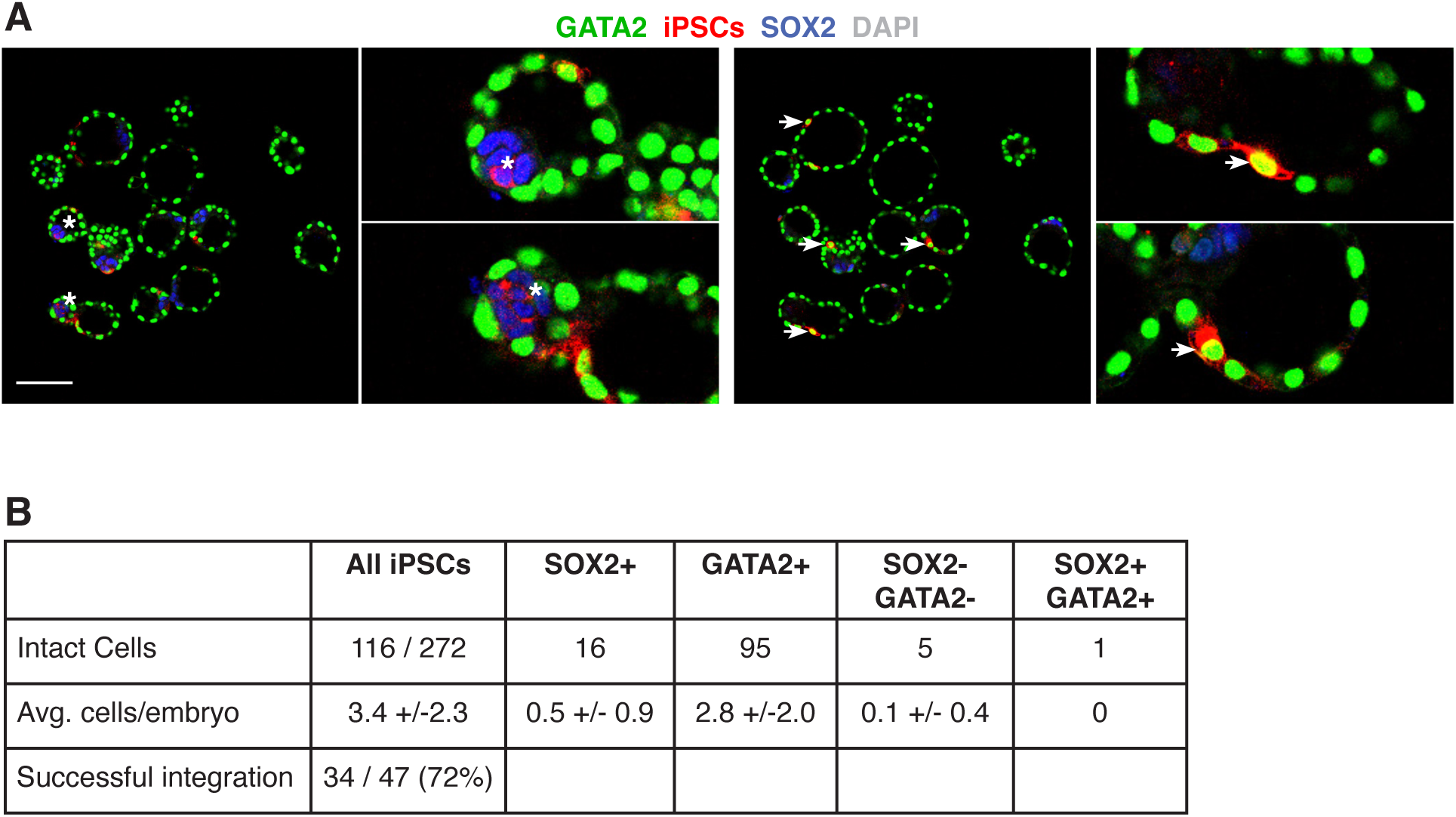
Dunnart iPSCs contribute to the inner cell mass and trophectoderm in early mouse blastocysts A) Immunofluorescence analysis of SOX2 (inner cell mass (ICM), blue), GATA2 (trophectoderm, green) expression in mouse blastocysts after injection at the 8-cell stage with dunnart iPSCs at the 8-cell stage, with iPSCS in red. Dunnart iPSCs integrate into the ICM and express SOX2 but not GATA2, and conversely iPSCs integrate into the trophectoderm and express GATA2 but not SOX2. B) Quantification of dunnart iPSC survival and contribution to two primary germ layers. Less than 1% of iPSCs express both GATA2 and SOX2.

### Embryoid bodies from diPSCs compartmentalise and express genes associated with differentiation after removal of exogenous reprogramming factors

As a second functional measure of pluripotency, C5/PLAX diPSCs were tested for their ability to differentiate into distinct lineages in an embryoid body (EB) formation assay. When cultured under low adherence conditions in the presence of doxycycline, large homogeneous aggregates of cells formed (Fig. 3A, 3 days (dox)). After removal of doxycycline and transfer to serum-containing medium, EBs grew in size and formed smooth outer surfaces and distinct internal structures (Fig. 3A, 9 days (no dox)). When layered onto ECM gel, EBs underwent significant organisation and compartmentalisation, with two distinct subsets of cellular organisation in different regions (Fig. 3B, asterisks at 18 days). To determine whether EBs derived from diPSCs exhibit heterogenous differentiation capacity by expressing markers associated with different early cell lineages, immunofluorescence was performed using antibodies specific for markers of the hypoblast (GATA6, Fig. 3C, red) and trophoblast (GATA2, Fig. 3C, green). Significant co-expression of nuclear localised GATA2 and GATA6 was observed in subpopulations of cells, with a further subpopulation expressing only GATA6 (Fig. 3C, arrows), and cells near the opposite pole of the EB being negative for both markers. This heterogeneity and organisation of cellular expression profiles was supported by assessment of the intermediate filament marker vimentin (VIM), which was highly expressed at one region (Supp. Fig. 7, asterisk) while being largely absent from the opposite region of EBs (Supp. Fig. 7, arrow). Even within the vimentin-positive pole, a mosaic pattern of expression was observed. Bulk RNAseq analysis further supported the differentiation potential of diPSCs through EB formation, with gene ontology analysis of the top 200 genes increased in EBs compared to iPSCs identifying several biological processes with established roles in cellular differentiation and tissue development, including “cellular development process”, “cell differentiation” and “anatomical structure development and morphogenesis” (Fig. 3D, and Supp. Table 5). Further, strong enrichment of biological processes involved in central nervous system development were also significantly enriched, including “neurogenesis”, “axon development” and “neuron projection”. These results support the notion that after maintenance of diPSCs under continued expression of exogenous reprogramming factors, the in-vitro differentiation capacity of diPSCs in EBs is maintained long-term after removal of exogenous factors.

### Mixed profile of epigenetic marks at pluripotency-associated genes in diPSCs

The ability of diPSCs to robustly differentiate in standard in-vitro assays indicates that activation of the endogenous marsupial pluripotency network was achieved in these cells, even in the presence of continued exogenous reprogramming factor expression. To determine whether the epigenetic landscape of diPSCs reflects this, active (H3K4me3 & H3K27ac) and repressive (H3K27me3) histone modifications were assessed in the presence of transgene expression using cleavage under targets and release using nuclease (CUT&RUN, Fig. 4) on an independent diPSC line. Widespread repression marks at somatic developmental genes like the HOXA cluster were identified, while a subset of pluripotency and associated factors showed activation marks, including *POU5F1, KLF4*, *KLF2, TFCP2L1* and *TBX3* (Fig. 4A, and Supp. Table 6-8). Repressive marks were also identified at a subset of pluripotency genes, most notably *SOX2* and *NANOG*, despite activation marks and high transcript levels of an important activator of *NANOG* in iPSCs, *SALL4*, and consistent with low levels of the NANOG target gene, *ESRRB* (Fig. 1D & Supp. Fig 8A). Of note, *NANOG* expression increased significantly when diPSCs were differentiated towards endoderm, which demonstrates that the NANOG locus is functionally responsive in diPSCs, and reflects the known role of NANOG in promoting endoderm formation via a non-cell autonomous mechanism^24,25^.

### Global analysis of active epigenetic marks shows strong enrichment of POU5F1, SOX2 and POU2F1 binding motifs in diPSCs

To infer global patterns in the regulation of transcription factor binding between iPSCs and fibroblasts, analysis of motif enrichment (AME) was performed on the CUT&RUN data. In diPSCs, enrichment of H3K4me3 active marks was found at binding regions for transcription factors involved in early embryonic development, particularly those governing pluripotency (POU5F1, SOX2, ZFP42), the first lineage segregation into trophoblast and inner cell mass (CDX2, GATA3, GATA6, SOX17), and very early cleavage/zygotic genome activation stages (DUX4, ARGFX, TPRX1, DPRX) (Fig. 4B, and Supp. Table 9). Notably, of the 350 transcription factor binding motifs identified as being enriched for H3K4me3 marks, 60–70% correspond to transcription factors whose roles are best-described as being involved in later embryonic development, including HOX genes, neural patterning genes (DLX, LHX, EMX, IRX, NKX, PHOX, VSX), mesodermal regulators (MEOX, PRRX), and tissue-specific developmental factors. We speculated that the presence of H3K4me3 marks at motifs for such transcription factors reflects sites that are poised for later activation, a common feature of pluripotent cells where developmental regulators carry active or bivalent chromatin marks before lineage commitment. In support of this, a major pluripotency regulator known to act in concert with POU5F1 is POU2F1^26^, which facilitates the maintenance of a poised state of transcription through an anti-repression mechanism^27^. We assessed H3K27ac motifs and revealed a strong enrichment for pluripotency, lineage specification, and early embryonic/developmental regulators, particularly those involved in the transition from the morula to the blastocyst and the earliest post-implantation events associated with trophoblast differentiation (Fig. 4B-C, and Supp. Table 10). Importantly, both *POU5F1* as well as *POU2F1*, alone or together with *SOX2*, represent 4/10 of the most significantly enriched transcription factor binding motifs containing H3K27Ac marks. Overall, pluripotency-associated gene enhancers are marked by H3K27AC, and this demonstrates these diPSCs, indicating a pluripotency-associated epigenetic circuit has been activated in these diPSCs. Further, transcription of POU2F1 was significantly increased in diPSCs compared with fibroblasts (Supp. Table 3). Altogether, epigenetic analysis demonstrated that widespread H3K4me3 and H3K27ac activation marks are present at key pluripotency and associated genes in diPSCs, though repressive marks are present at a subset of these genes. Further, motif enrichment showed that a POU5F1- and SOX2-driven pluripotency network is active in diPSCs and indicates a conserved role for POU2F1 in priming the dunnart genome for pluripotency exit.

### diPSCs contribute to inner cell mass and trophoblast lineages after injection into mouse embryos

With epigenetic analysis of activation marks revealing enrichment of pluripotency- and trophoblast-associated motifs, it was hypothesised that diPSCs can contribute to both embryonic and extraembryonic lineages upon differentiation after removal of exogenous factors. As methods for dunnart embryo injection have yet to be developed, diPSCs were injected into 8-cell stage mouse embryos to test this hypothesis. After 48-72 hours, iPSC-derived cells survived and attained morphological features similar to the adjacent mouse cells (Fig. 5A, red). Quantitative analysis showed that 72% of embryos injected had at least one successful diPSC engraftment, representing 43% of all cells injected. (Fig. 5B). When embryos were co-immunolabelled for a marker of the inner cell mass, SOX2 (Fig 5A, blue), and the trophoblast marker GATA2 (green), results showed that diPSCs contributed to both the inner cell mass and trophoblast. Importantly, aside from a single diPSC expressing both SOX2 and GATA2, SOX2-positive diPSCs of the inner cell mass did not express GATA2, while GATA2-positive diPSCs of the trophoblast did not express SOX2. Together this demonstrates that diPSCs survive and integrate into early mouse embryos, can respond to external differentiation cues, and contribute to the earliest mouse embryonic and extraembryonic lineages.

## Discussion

Our study successfully generated and maintained diPSCs without the need for removal of exogenous factors. Importantly, our diPSCs exhibited the necessary properties suggesting suitability for downstream marsupial conservation pipelines, including the demonstration that continued expression of exogenous reprogramming factors does not impede the capacity of cells to differentiate in standard in-vitro assays of pluripotency after removal of the exogenous factors. This finding is important for the field of next-generation conservation as it demonstrates that achieving transgene-free stability of iPSCs in relevant species is not essential to establish tools for the application of iPSCs in conservation-relevant species, potentially expediting pipelines and more efficiently applying investments that will ultimately improve outcomes for species rescue.

The diPSCs developed in this study exhibit a pluripotency phenotype sharing many features with eutherian iPSCs, including the capacity to differentiate into all three embryonic germ lineages, express trophoblast and primitive endoderm markers in EB differentiation assays, and contribute to both embryonic and extraembryonic lineages in early mouse blastocysts. Together, our data provides strong support that the large evolutionary distances between marsupials and eutherians and the unique complexities of marsupial pluripotency (possessing both *POU5F1* and *POU5F3*) are not major barriers to success in applying generalised approaches to species rescue through somatic cell reprogramming. While it remains to be seen whether such iPSCs can integrate in dunnart embryos, the use of an inducible system to generate iPSCs from somatic cells facilitates the discovery of specific culture conditions able to facilitate transgene independence. Further, an inducible approach enables convenient application of transient reprogramming factor expression to rejuvenate the epigenetic status of starting material. This is particularly relevant in the context of threatened and endangered species where starting material is more likely to be of relatively low quality due derivation from aged animals, from a limited number of individuals, or from biobanks where material may have been collected in circumstances that compromise cellular integrity. Even if iPSCs derived from lines exposed to continued exogenous reprogramming factor expression are found to be less suitable for application in generating live animals, they still provide an excellent tool to study pluripotency in each at-risk species, facilitating the application of next-generation conservation technologies in the future.

The diPSCs generated in this study exhibit many of the transcriptional and epigenetic hallmarks of pluripotency, including expression of core pluripotency factors, widespread repression of somatic genes, and activation of pluripotency-associated genes^28^. However, the overall transcriptomic and epigenomic profile of pluripotency-associated genes in our diPSCs was mixed, with some genes highly active (e.g. *POU5F1*, *TFCP2L1*, *KLF2*), some repressed (e.g. *NANOG*, *SOX2*) and some bivalent marks associated with only small increases in expression compared with fibroblasts (e.g. *PRDM14*). In the case of *SOX2* and *NANOG*, this was associated with repression of endogenous expression. It was not possible to distinguish whether this is a consequence of persistent expression of exogenous pluripotency factors resulting in repression of a subset of endogenous factors, or due to suboptimal culture conditions or an incomplete reprogramming phenotype resulting in somatic epigenetic memory^29^. Nonetheless, the observed robust differentiation potential suggests that the diPSCs generated in this study are truly pluripotent. Further, several response elements of genes involved in regulating differentiation of the early embryo were enriched with active promoters/enhancers, including those associated with the inner cell mass, trophoblast and germ cell lineages, as well as those known to regulate substates of pluripotency (*TFAP2C, TFAP2A*, *KLF2* and *KLF4* ^30, 31, 32, 33, 34^). A more definitive level of evidence would require dunnart embryo injections to assess the capacity of diPSCs to contribute to all germ layers in this more relevant biological context. Such methods have not been established for a marsupial, although significant investments are currently being made towards this end, along with additional assisted reproduction technologies for marsupials such as genome editing *in ovo*, embryo transfer, and spermatogonial stem cell enrichment, culture and transplantation.

It is worth highlighting the absence of activation marks at, and transcriptional activity of, *NANOG* & *ESRRB* in stable diPSCs. This observation is inconsistent with previous studies which reported *NANOG* expression in other marsupial iPSCs^1, 2^, and in one case *NANOG* was required for successful reprogramming^2^. Additionally, *NANOG* expression was also reported in iPSCs of a monotreme species^7^. The present study demonstrated that significant endogenous *NANOG* expression does occur in partially reprogrammed dunnart cells and is highly induced during early endoderm differentiation of diPSCs, indicating that exogenous *NANOG* is not required for successful reprogramming of dunnart somatic cells, as exogenous reprogramming factors are able to activate endogenous *NANOG* expression at levels sufficient for cells to navigate the reprogramming process. Continued expression of *NANOG* and its downstream target, *ESRRB*, were not required to maintain pluripotency in diPSCs, at least in the context of continued exogenous reprogramming factor expression. In the opossum study, when compared to samples from different stages of early embryonic development iPSCs most resembled the epiblast/hypoblast stage, while in uncommitted cells NANOG expression was very low^2^. It is tempting to speculate that the absence of active NANOG in diPSCs may be due to the cells occupying an earlier, more naïve state of pluripotency under the specific reprogramming and culture conditions of the present study. It is also possible that similar to findings in the opossum and Tasmanian devil, ESRRB is not as heavily engaged in maintaining pluripotency in diPSCs. Further, published data suggests that pluripotency is principally maintained by an OCT4-SOX2 regulatory module, while NANOG and ESRRB operate a partially distinct network involved in regulating the timing of differentiation and stability of the pluripotent state^35, 36^. When combined, results of previous studies and the present study may suggest a bias towards the former module in marsupials. However, because no marsupial iPSCs have successfully been used to generate chimeras to functionally assess the lack of ESRRB expression, it is equally possible that all marsupial iPSCs generated to date occupy a reprogrammed state not associated with full pluripotency. This highlights the immediate need for the development of ART to progress the field of stem cell biology for marsupial conservation.

Altogether, this study demonstrates that diPSCs can be generated and maintained for application in downstream next-generation species rescue efforts. Where culture conditions are not sufficiently refined to enable the maintenance of an exogenous reprogramming factor-free state of pluripotency, continued exogenous expression does not impede application of the resulting iPSCs. These observations provide a path for the use of iPSCs in the context of species rescue, where cellular resources may be limited and species-specific culture conditions are unknown.

## Acknowledgements

The authors wish to acknowledge the contributions of Jinny Chang and Paul McMillan from The University of Melbourne Biological Optical Microscopy Platform, Naomi Tappe for technical assistance in embryo injections, and both the Center for Genome Innovation and Computational Biology Core in the UConn Institute for Systems Genomics for sequencing and HPC support, respectively.

