## Supplementary Figures for "Rapid generation of marsupial induced pluripotent stem cells suitable for use in next-generation conservation"

### Supp. Fig. 1

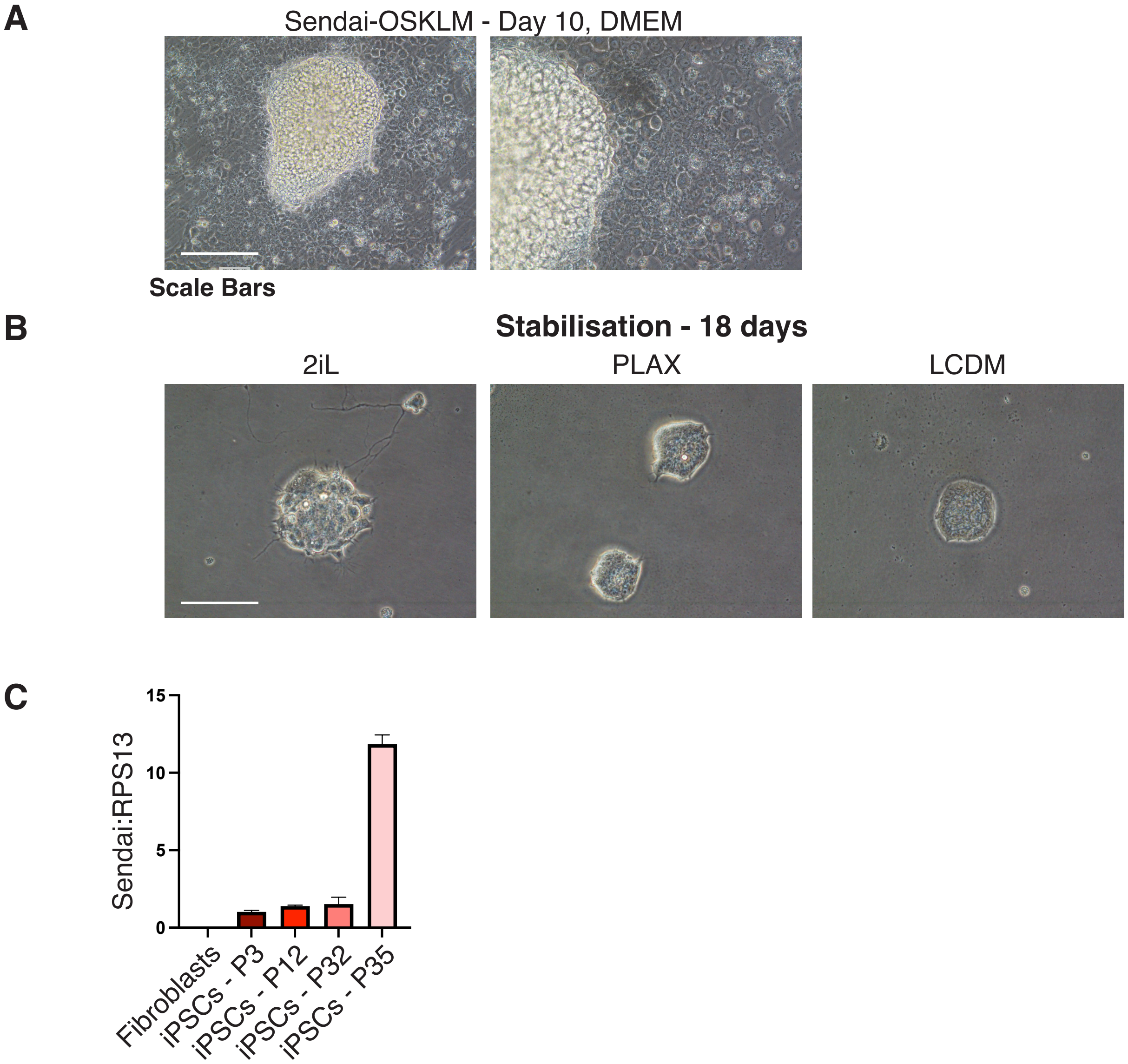


**Supp. Fig. 1 - Persistence of sendai virus-driven reprogrammming factor expression in diPSCs after long-term passage** A) Morphological changes in dunnart fibroblasts after infections with Sendai virus-driven expression of reprogramming factors. B) Morphology of colonies picked from A) and cultured under different conditions. C) qPCR of sendai virus RNA over passage in diPSCs. (Bar = 100 µm).

**Supp. Fig. 2**

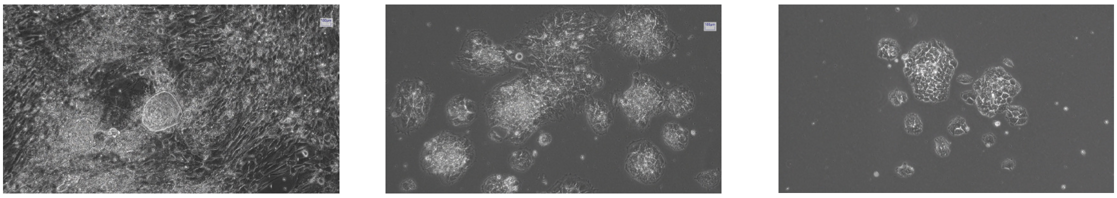


**Supp. Fig. 2 - RNA-based reprogramming failed to generate stable dunnart iPSCs.** Morphological changes in dunnart fibroblasts after transfection with RNA coding for reprogramming factors.

**Supp. Fig. 3**

**A**

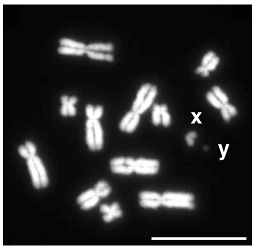


**B**

| iPSC Line | Proportion of normal karyotypes | Notes on abnormalities |
|---|---|---|
| scCDL24D (P35) | 16/19 | 3 with additional chromosome |
| scCDL24F (P32) | 32/36 | 1 tetraploid, 3 with additional chromosome |

**C**

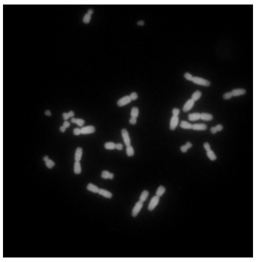

**Supp. Fig. 3 - Karyotype analysis of dunnart iPSCs after long-term passage.** A) Example of dunnart iPSC with a normal karyotype (Bar = 25 µm). B) Quantification of the proportion of normal and abnormal karyotypes in dunnart iPSCs after long-term passage. C) Example of an abnormal karyotype identified in a dunnart iPSC after long-term passage (Bar = 50 µm).

**Supp. Fig. 4**

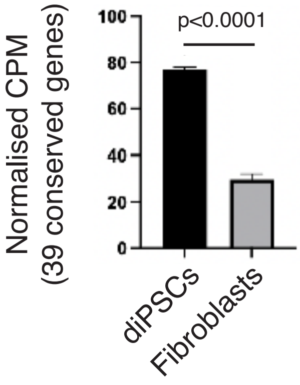


**Supp. Fig. 5**

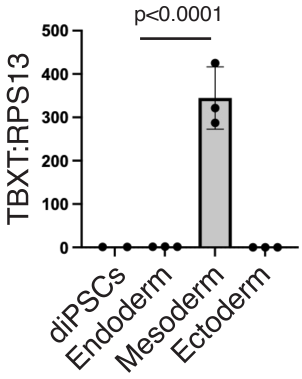


**Supp. Fig. 4 - Normalised expression of 39 genes conserved in iPSCS from diverse species (mouse, pig, platypus, chicken) as determined by RNAseq.**

**Supp. Fig. 5 - Expression of mesoderm marker, TBXT, is restricted to mesoderm-differentiated iPSCs after extended trilineage differentiation, as determined by RT-qPCR.**

### Supp. Fig. 6

#### Endoderm

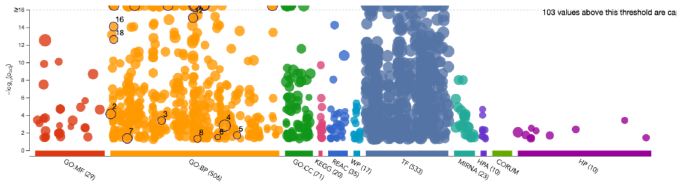


| ID | Source | Term ID | Term Name | $p_{adj}$ (query_1) |
|---|---|---|---|---|
| 1 | GO:BP | GO:0048856 | anatomical structure development | $2.786\times10^{-51}$ |
| 9 | GO:BP | GO:0035295 | tube development | $1.002\times10^{-32}$ |
| 10 | GO:BP | GO:2000026 | regulation of multicellular organismal developm... | $6.953\times10^{-30}$ |
| 15 | GO:BP | GO:0060429 | epithelium development | $1.594\times10^{-27}$ |
| 14 | GO:BP | GO:0001944 | vasculature development | $2.264\times10^{-24}$ |
| 13 | GO:BP | GO:0048762 | mesenchymal cell differentiation | $2.817\times10^{-19}$ |
| 17 | GO:BP | GO:0060485 | mesenchyme development | $9.282\times10^{-19}$ |
| 11 | GO:BP | GO:0009790 | embryo development | $1.597\times10^{-17}$ |
| 12 | GO:BP | GO:0048598 | embryonic morphogenesis | $7.485\times10^{-16}$ |
| 16 | GO:BP | GO:0001822 | kidney development | $7.508\times10^{-15}$ |
| 18 | GO:BP | GO:0001837 | epithelial to mesenchymal transition | $2.251\times10^{-13}$ |
| 2 | GO:BP | GO:0000122 | negative regulation of transcription by RNA pol... | $6.819\times10^{-5}$ |
| 3 | GO:BP | GO:0032963 | collagen metabolic process | $3.778\times10^{-4}$ |
| 4 | GO:BP | GO:0080090 | regulation of primary metabolic process | $1.428\times10^{-3}$ |

#### Mesoderm

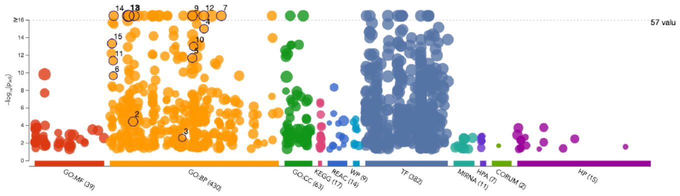


| ID | Source | Term ID | Term Name | $p_{adj}$ (query_1) |
|---|---|---|---|---|
| 1 | GO:BP | GO:0007275 | multicellular organism development | $9.675\times10^{-47}$ |
| 13 | GO:BP | GO:0007399 | nervous system development | $1.380\times10^{-29}$ |
| 12 | GO:BP | GO:0060429 | epithelium development | $4.156\times10^{-27}$ |
| 7 | GO:BP | GO:0072359 | circulatory system development | $7.022\times10^{-22}$ |
| 8 | GO:BP | GO:0009790 | embryo development | $1.603\times10^{-20}$ |
| 14 | GO:BP | GO:0001944 | vasculature development | $4.436\times10^{-17}$ |
| 9 | GO:BP | GO:0048598 | embryonic morphogenesis | $8.475\times10^{-17}$ |
| 4 | GO:BP | GO:0060485 | mesenchyme development | $9.624\times10^{-16}$ |
| 15 | GO:BP | GO:0001501 | skeletal system development | $4.607\times10^{-14}$ |
| 10 | GO:BP | GO:0048762 | mesenchymal cell differentiation | $9.904\times10^{-14}$ |
| 5 | GO:BP | GO:0048568 | embryonic organ development | $2.032\times10^{-12}$ |
| 11 | GO:BP | GO:0001822 | kidney development | $4.290\times10^{-12}$ |
| 6 | GO:BP | GO:0001837 | epithelial to mesenchymal transition | $2.172\times10^{-10}$ |
| 2 | GO:BP | GO:0009611 | response to wounding | $3.580\times10^{-5}$ |
| 3 | GO:BP | GO:0045109 | intermediate filament organization | $2.570\times10^{-3}$ |

#### Ectoderm

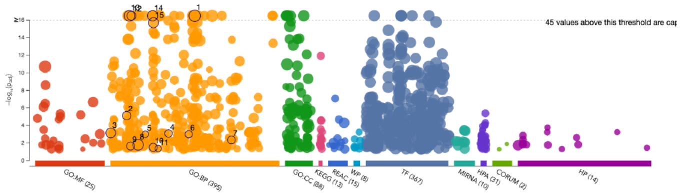


| ID | Source | Term ID | Term Name | $p_{adj}$ (query_1) |
|---|---|---|---|---|
| 1 | GO:BP | GO:0048856 | anatomical structure development | $1.070\times10^{-40}$ |
| 12 | GO:BP | GO:0008544 | epidermis development | $1.278\times10^{-19}$ |
| 13 | GO:BP | GO:0007399 | nervous system development | $3.641\times10^{-19}$ |
| 14 | GO:BP | GO:0022008 | neurogenesis | $8.456\times10^{-17}$ |
| 15 | GO:BP | GO:0030182 | neuron differentiation | $1.965\times10^{-16}$ |
| 2 | GO:BP | GO:0006814 | sodium ion transport | $7.567\times10^{-6}$ |
| 3 | GO:BP | GO:0000122 | negative regulation of transcription by RNA pol... | $7.069\times10^{-4}$ |
| 4 | GO:BP | GO:0035418 | protein localization to synapse | $8.149\times10^{-4}$ |
| 5 | GO:BP | GO:0018149 | peptide cross-linking | $9.858\times10^{-4}$ |
| 6 | GO:BP | GO:0046513 | ceramide biosynthetic process | $9.938\times10^{-4}$ |
| 7 | GO:BP | GO:0098659 | inorganic cation import across plasma membra... | $4.466\times10^{-3}$ |
| 8 | GO:BP | GO:0010557 | positive regulation of macromolecule biosynthe... | $1.596\times10^{-2}$ |
| 9 | GO:BP | GO:0007612 | learning | $2.197\times10^{-2}$ |
| 10 | GO:BP | GO:0030010 | establishment of cell polarity | $3.130\times10^{-2}$ |
| 11 | GO:BP | GO:0032252 | secretory granule localization | $4.183\times10^{-2}$ |

**Supp. Fig. 6 - Gene ontology analysis of RNAseq data from individual trilineage differentiation samples.**

### Supp. Fig. 7

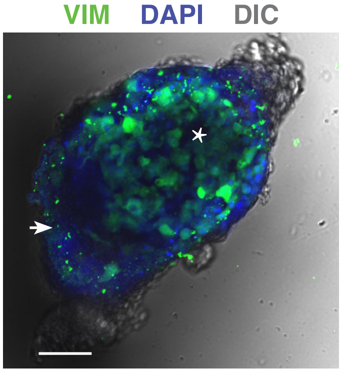


**Supp. Fig. 7 - Vimentin Immunostaining in dunnart iPSC-derived embryoid bodies.** After transfer to ECM substrate, distinct compartments emerge in embryoid bodies, as illustrated by low/ absent vimentin at one region (arrow), and widespread but variable expression at the opposing region (asterisk). Bar = 100 µm.

### Supp. Fig. 8

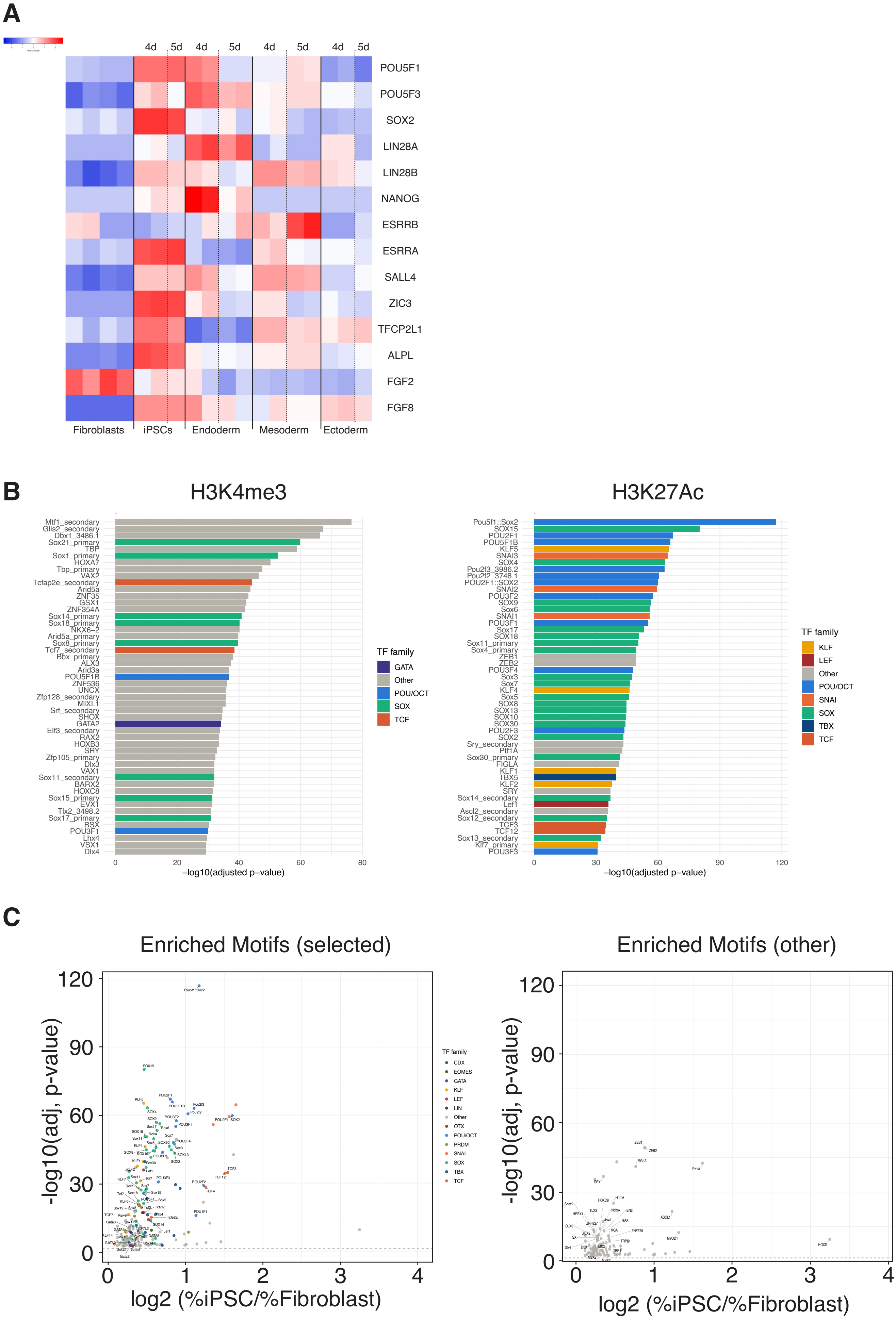
A
4d
5d
4d
5d
4d
5d
4d
5d
POU5F1
POU5F3
SOX2
LIN28A
LIN28B
NANOG
ESRRB
ESRRA
SALL4
ZIC3
TFCP2L1
ALPL
FGF2
FGF8
Fibroblasts
iPSCs
Endoderm
Mesoderm
Ectoderm
B
H3K4me3
H3K27Ac
TF family
GATA
Other
POU/OCT
SOX
TCF
KLF
LEF
SNAI
TBX
−log10(adjusted p−value)
C
Enriched Motifs (selected)
Enriched Motifs (other)
-log10(adj, p-value)
log2 (%iPSC/%Fibroblast)
TF family
CDX
EOMES
GATA
KLF
LEF
LIN
Other
OTX
POU/OCT
PRDM
SNAI
SOX
TBX
TCF
